# Phosphoproteomics profiling of the monocytes uncovers the mechanosensory pathways regulating monocytes responses to pathologic level of shear stress

**DOI:** 10.64898/2026.10.06.757199

**Authors:** Austin Lai, Chanly Chheang, Haoyun Fang, Habiba Danish, Yasmin Mirzaalikhan, Nalin Dayawansa, Anna Watson, Jonathan Noonan, Ching-Seng Ang, David Greening, Karlheinz Peter, Sara Baratchi

**Affiliations:** Baker Heart and Diabetes Institute, Melbourne, Victoria, Australia; School of Health and Biomedical Sciences, RMIT University, Melbourne, Victoria, Australia; Baker Department of Cardiometabolic Health, University of Melbourne, Melbourne, Victoria, Australia; Bio21 Molecular Science and Biotechnology Institute, The University of Melbourne, Melbourne, Victoria, Australia; Baker Department of Cardiovascular Research Translation and Implementation, La Trobe University, Bundoora, Victoria, Australia

## Abstract

**Background:** Calcific aortic valve disease (CAVD) is the most common valvular heart disease in older adults and currently has no approved pharmacological therapy to prevent or slow disease progression. Aortic valve replacement, either surgically or by transcatheter aortic valve implantation (TAVI), therefore remains the only effective treatment for severe CAVD. Monocytes play an important role in driving inflammation and disease progression in CAVD. As the disease progresses and the aortic valve becomes increasingly stenotic, circulating monocytes passing through the narrowed valve are repeatedly exposed to pathological haemodynamic forces, including markedly elevated shear stress, before their recruitment into the valve tissue. However, how exposure to these mechanical forces reprograms circulating monocytes influences their subsequent responses within the valve remain poorly understood.

**Methods and Results:** In this study, we used an engineered model of aortic valve stenosis combined with phosphoproteomic profiling to investigate how prolonged exposure to pathological shear stress reprograms circulating monocytes and influences their subsequent phenotype and function following recruitment into the valve matrix. We further used TAVI as a unique clinical model to determine whether restoration of physiological haemodynamics in patients with severe aortic stenosis reverses the shear-induced monocyte phenotype observed in our experimental model. We found that prolonged exposure of monocytes to pathological shear stress induced extensive remodelling of the proteomic and phosphoproteomic landscape, with enrichment of pathways associated with cellular metabolism, cytoskeletal organisation, and inflammatory signalling. Kinase activity analysis further identified increased protein kinase C (PKC) activity accompanied by suppression of AKT signalling. Consistent with these molecular changes, pathological shear stress increased actin polymerisation, mitochondrial membrane potential, intracellular calcium signalling, and PKC activity. Using a bioengineered extracellular matrix model that recapitulates the glycosaminoglycan-rich microenvironment of the stenotic aortic valve we found that prior exposure to pathological shear stress altered monocyte-to-macrophage differentiation, reducing oxidised LDL uptake and foam-cell formation while promoting co-expression of CD80 and CD163, indicative of a mixed macrophage phenotype. Finally, using a myeloid-specific Piezo1 knockout mouse model, we identified Piezo1 as a key mechanosensor linking pathological shear stress to PKC activation, mitochondrial responses, and persistent changes in monocyte phenotype following migration into the extracellular matrix.

**Conclusions:** These findings establish a mechanistic link between abnormal valve haemodynamics, metabolic and inflammatory reprogramming of circulating monocytes, extracellular matrix remodelling, and immune dysregulation during CAVD progression, highlighting monocyte mechanotransduction and downstream PKC/Piezo1 signalling as potential therapeutic targets.

## Introduction

Calcified aortic valve disease (CAVD) is the most prevalent type of valvular heart disease in developed countries, with a significant impact on the elderly population. The prevalence of CAVD increases with age, affecting approximately 0.2% of individuals aged 50-59 and rising sharply to around 10% in those aged 80-89 ^1, 2^. The disease is characterized by progressive calcification and stiffening of the aortic valve, which leads to a narrowing of the valve opening, a condition known as aortic stenosis. This narrowing impedes blood flow from the heart into the aorta, causing increased workload on the heart and potentially leading to heart failure if left untreated ^3^.

The standard treatment for severe CAVD is either surgical aortic valve replacement (SAVR) or transcatheter aortic valve implantation (TAVI), a minimally invasive procedure in which a prosthetic valve is delivered via a catheter to replace the diseased valve ^3^. Currently, there are no approved pharmacological treatments specifically targeting the progression of aortic stenosis, highlighting a significant gap in the therapeutic landscape. This lack of pharmacotherapy underscores the importance of advancing our understanding of the disease at a molecular level, which could lead to the development of novel drugs or therapeutic strategies. CAVD initiation and progression are driven by chronic inflammation and altered valvular haemodynamics, which expose circulating blood cells to pathological mechanical forces ^4^. As blood recirculates continuously, inflammatory cells especially larger cells like monocytes experience thousands of episodes of supraphysiological shear stress during their lifespan as they passed through the narrowed and calcified aortic valve^5^. Previous studies by our group have established a link between these recurrent pathological mechanical stimuli and activation of inflammatory signalling pathways in circulating blood cells such as monocytes ^6^ and neutrophils ^7^. However, whether shear stress–induced responses persist after monocytes leave the circulation and differentiate into macrophages within the valvular microenvironment remains unknown. Understanding which shear stress-induced responses persist in monocyte-derived macrophages is essential for identifying specific regulatory pathways that could be therapeutically targeted to modulate immune responses in CAVD.

One of the earliest events in mechanotransduction is the rapid induction of post-translational modifications (PTMs), particularly phosphorylation, which convert mechanical stimuli into intracellular biochemical signals ^8, 9^. This mechanism has been extensively studied in endothelial cells, where phosphorylation regulates multiple aspects of mechanosensitive signalling ^10–13^. In monocytes, however, the role of phosphorylation in mechanotransduction remains less well understood. Nevertheless, such protein modifications, particularly phosphorylation, play a critical role in regulating monocyte activation, metabolic reprogramming, and inflammatory phenotype during CAVD ^14^. For example, monocytes isolated from patients with aortic stenosis exhibit altered phosphorylation profiles, characterised by increased phosphorylation of metabolic regulators such as pyruvate dehydrogenase kinase 4 (PDK4) and reduced phosphorylation of immunomodulatory kinases, including inhibitor of nuclear factor kappa-B kinase subunit epsilon (IKBKE) ^15^. However, the upstream mechanosensors responsible for initiating these phosphorylation-dependent signalling events in monocytes remain poorly defined.

Piezo1 has emerged as a key mechanosensitive ion channel regulating immune cell responses to mechanical forces. Structurally, Piezo1 forms a homotrimeric, three-bladed propeller-like non-selective cation channel that primarily conducts Na^+^, K^+^, and Ca^2+^, enabling the conversion of mechanical stimuli into intracellular biochemical signals ^16–20^. Piezo1 is highly expressed in monocytes and macrophages, where it is activated by substrate stiffness, shear stress, cyclic stretch, and chemical agonists ^17, 21–23^. In macrophages, Piezo1-mediated Ca^2+^ influx promotes NF-κB-dependent pro-inflammatory signalling while suppressing anti-inflammatory polarization, thereby regulating macrophage phenotype in response to matrix stiffness ^24^. Importantly, accumulating evidence demonstrates that Piezo1 is also a critical mediator of shear stress-induced monocyte activation. High shear stress activates Piezo1, leading to increased expression of inflammatory markers, including CD11b, IL-6 and IFN-β1, enhanced oxidised LDL uptake and phagocytosis, responses that are recapitulated in monocytes isolated from patients with aortic stenosis and attenuated following TAVI ^5^. Furthermore, Piezo1 has been shown to regulate shear-induced inflammasome activation through the IKKβ–NLRP3 pathway in macrophages ^25^, while also mediating inflammatory signalling in response to cyclic mechanical loading through Ca^2+^-dependent activation of c-JUN/AP-1 and HIF-1α pathways ^21, 26^. Collectively, these findings establish Piezo1 as a central regulator of immune mechanotransduction and suggest that it may represent an attractive therapeutic target for limiting pathological inflammatory responses induced by abnormal haemodynamic forces ^27, 28^.

## Materials and Methods

### Cell culture

THP-1 cells were cultured in complete RPMI-1640 medium containing 10% fetal bovine serum, 1% penicillin-streptomycin and 10 mM HEPES (all obtained from Thermo Fisher Scientific). The same media formulation was used for isolated human and mouse monocytes.

### Animals

To generate myeloid-specific Piezo1 knockout (Piezo1 KO) *Piezo1*^Δ*lysM*^ mice, *Piezo1^flox/flox^* (Piezo1tm2.1Apat, Jackson Laboratories stock no: 029213) were first bred with *LysM^Cre/Cre^* (B6.129P2-Lyz2tm1(cre)Ifo/J, Jackson Laboratories stock no: 004781) mice to generate *Piezo1^flox/+^ LysM^Cre/+^* heterozygotes. Further breeding of the heterozygotes with *Piezo1^flox/flox^*generated *Piezo1*^Δ*lysM*^ mice. Animals of 12-18 weeks were used in this study. Animal experiments were approved by the Alfred Health Research Ethics Committee. The knockdown of Piezo1 was confirmed using qPCR and response to Yoda1 *in vitro*.

### Monocyte isolation

This study used human monocytes isolated from blood of human donors with approval from the Alfred Hospital Research Ethic Committee. Blood (20-50 ml) was obtained in citrate 3.2%-coated vacutainers (S-Monovette® 8.2ml 9NC, Sarstedt) from healthy donors. Ficoll® Paque Plus (Sigma-Aldrich) and Histopaque-1119 (Sigma-Aldrich) were used to generate a density gradient in which the blood was carefully layered on top followed by centrifugation to separate peripheral blood mononuclear cells (PBMCs) from blood, as described previously^29^. The PBMC pellet was suspended in RPMI-1640 medium and subjected to shear stress and further assessed in downstream applications described below. For mice experiments, femurs were obtained from mice and submerged in complete RPMI medium. Bone marrow cells from femurs were flushed with complete RPMI medium and bone-marrow-derived monocytes were isolated using a mouse monocyte isolation kit (Miltenyi Biotec).

## Ethic statements

The study protocol was approved by the Alfred Human Research Ethics Committee (Project No. 121/22), and written informed consent was obtained from all participants prior to enrolment. Animal experiments were conducted with approval from the Alfred Medical Research and Education Precinct Animal Ethics Committee (Project No. 8217). Recruitment of patients undergoing TAVI was approved by the Alfred Hospital Human Research Ethics Committee (Project No. 560/1). All human studies were performed in accordance with the principles of the Declaration of Helsinki.

### *In vitro* generation of shear stress

Prior to experimentation, µ-Slide VI 0.1 flow chambers (ibidi) and collection tubes were coated with 5% bovine serum albumin (BSA) for at least 1 h at 37°C to prevent cell adhesion. Tubing (0.09-inch ID; EZkem) was connected to a LabV1 peristaltic pump (Shenchen), sterilised using ultraviolet light and ethanol, and subsequently coated with 5% BSA. Following isolation, PBMCs in suspension were perfused through the system and exposed to flow rates of 1 ml/min (≈80 dyne/cm², low shear stress (LSS)) or 5 ml/min (≈400 dyne/cm², high shear stress (HSS)) for 10 min, 30 min, 6 h, or 24 h at 37°C. To investigate the role of PKC, cells were pre-treated with the PKC inhibitors Rottlerin (2 µM; Calbiochem) or Calphostin C (10 nM; Calbiochem) for 30 min, AKT inhibitor GSK690693 (100 nM; Selleck Chemicals), cyclin dependent kinase inhibitors Flavopiridol (10 nM; MedChemExpress) or Ro-3306 selective for cyclin-dependent kinase 1 (5 µM; MedChemExpress), or casein kinase 2 inhibitor Ellagic acid (5 µM; MedChemExpress) before shear stress exposure, with inhibitors maintained throughout the 6 h shear stress period. Following shear stress, PBMCs were seeded into 8-well chamber slides (ibidi) for 10–15 min to allow monocyte adhesion, after which non-adherent cells were removed by gentle washing. For proteomic and phosphoproteomic analyses, THP-1 cells were collected following shear stress exposure. Adherent monocytes were subsequently processed for downstream assessments via encapsulation within collagen hydrogels, and used for gene expression and immunocytochemical analyses as described below. For intracellular calcium ([Ca²]) measurements, Fluo-4 AM (1 µM; Invitrogen) was added to the medium during shear stress exposure.

### Proteomics and phosphoproteomics sample preparation

Cells were washed twice with pre-warmed PBS to remove residual serum and lysed directly in pre-heated guanidine hydrochloride (GdnHCl) lysis buffer containing 6 M GdnHCl, 100 mM Tris (pH 8.5), 5 mM tris(2-carboxyethyl)phosphine (TCEP) and 10 mM chloroacetamide. Lysates were heated at 95°C for 10 min in the dark and mechanically sheared to disrupt genomic DNA. Samples were centrifuged at 13,000 rpm for 5 min at 4°C, and the supernatants were collected.

Approximately 500 µg of protein per sample was precipitated by addition of five volumes of ice-cold acetone/methanol and incubated overnight at −20°C. Samples were centrifuged at 13,000 rpm for 10 min at 4°C, and the resulting protein pellets were washed once with ice-cold acetone. Pellets were completely resuspended in 100 µL of 6 M GdnHCl by alternating vortexing and sonication and subsequently diluted with 500 µL of 50 mM Tris-HCl (pH 8.0) to reduce the final GdnHCl concentration to approximately 1 M. Protein concentration was determined using a Micro BCA Protein Assay, and samples were normalised to equal protein concentrations and volumes using 50 mM Tris-HCl (pH 8.0).

Proteins were digested with Trypsin/Lys-C at an enzyme-to-protein ratio of 1:50 and incubated overnight at 37°C with agitation. Following digestion, samples were acidified to 1% (v/v) formic acid and subjected to solid-phase extraction (SPE) using Oasis HLB cartridges (60 mg; Waters). Cartridges were conditioned with 80% acetonitrile (ACN) containing 0.1% trifluoroacetic acid (TFA) and equilibrated with 0.1% TFA. Samples were loaded onto the cartridges, washed with 0.1% TFA and eluted with 80% ACN containing 0.1% TFA. Approximately 20 µg of peptide from each sample was reserved for global proteomic analysis, while the remaining peptide material was used for phosphopeptide enrichment. Samples were partially dried by vacuum centrifugation to remove ACN and subsequently lyophilised.

### Phosphopeptide enrichment

Phosphopeptides were enriched using titanium dioxide (TiO□) beads with a lactic acid-based enrichment protocol. Lyophilised peptides were resuspended in TiO□ loading buffer containing 2 M lactic acid, 5% TFA and 50% ACN. TiO□ beads were equilibrated sequentially with washing buffer (50% ACN, 5% TFA) and loading buffer. Peptides were incubated with TiO□ beads at a TiO□-to-peptide ratio of 6:1 for 1 h with agitation.

Following incubation, the peptide–TiO□ suspensions were transferred to C8-packed tips, and the flow-through was passed through the TiO□ matrix a second time to maximise phosphopeptide binding. The TiO□ matrix was washed twice with loading buffer followed by three washes with 50% ACN containing 5% TFA. Bound phosphopeptides were eluted three times with 1% (v/v) NH□OH (pH 11.3), followed by a final elution with 30% ACN. Eluted phosphopeptides were immediately acidified with formic acid prior to LC–MS/MS analysis.

### LC-MSMS

DIA-MS/MS was carried out on an Orbitrap Eclipse (Thermofisher Scientific) coupled to a Vanquish NEO (Thermofisher Scientific) nanoLC system. The LC system was equipped with an Acclaim Pepmap nano-trap column (Dinoex-C18, 100 Å, 75 µm x 2 cm) and an Acclaim Pepmap RSLC analytical column (Dinoex-C18, 100 Å, 75 µm x 50 cm). The tryptic peptides were injected to the enrichment column at an isocratic flow of 5 µL/min of 2% v/v CH_3_CN containing 0.1% v/v formic acid for 5 min applied before the enrichment column was switched in-line with the analytical column. The eluents were 5% DMSO in 0.1% v/v formic acid (solvent A) and 5% DMSO in 100% v/v CH_3_CN and 0.1% v/v formic acid (solvent B). The flow gradient was (i) 0-6min at 3% B, ii) 6-7min, 3-4% (ii) 7-82 min, 4-25% B (iii) 82-86min 25-40% B (iv) 86-87min, 40-80% B (v) 87-90min, 80-80% B (vi) 90-91min, 80-3% and equilibrated at 3% B for 10 minutes before the next sample injection. For DIA experiments, full MS resolutions were set to 120,000 at m/z 200 and scanning from 350-1400m/z in the profile mode. Full MS AGC target was 250% with an IT of 50 ms. AGC target value for fragment spectra was set at 2000%. 50 windows of 13.7 Da were used with an overlap of 1 Da. Resolution was set to 30,000 and maximum IT to 55 ms. Normalized collision energy was set at 30%. All data were acquired in centroid mode using positive polarity.

### Bioinformatics analysis of global and phosphoproteomics data

Raw DIA-MS/MS files were processed using the direct DIA analysis workflow in **Spectronaut** (v17.4.230317.55965) against the reviewed **UniProt Homo sapiens** database (downloaded April 2023). Trypsin specificity was specified with up to two missed cleavages. Carbamidomethylation of cysteine was defined as a fixed modification, while protein N-terminal acetylation and methionine oxidation were included as variable modifications. For phosphoproteomic analyses, phosphorylation of serine, threonine and tyrosine was included as an additional variable modification. Protein, peptide and peptide-spectrum match false discovery rates were controlled at 1%. DIA identifications were filtered using peptide posterior error probability and Q-value thresholds of 0.01. Quantification was performed at the MS2 level, with precursor filtering based on Q-values and cross-run normalisation performed using the automatic setting in Spectronaut.

Perseus (v2.0.7.0) bioinformatic software was used for preliminary global proteomics and phosphoprotoemics data analysis. Here, protein groups and phosphosites were log2-transformed with minimum 2 out of 3 valid values per group included for downstream analysis. Missing values were replaced by a left-centred normal distribution using Perseus default imputation settings (width: 0.3, down shift: 1.8). For differential analysis, the multi-group comparison was conducted using one-way ANOVA with permutation-based FDR reported and post-hoc analysis enabled. For pairwise comparison, student’s t-test was conducted with permutation-based FDR enabled. Downstream bioinformatics analysis and data visualization were conducted using R programming language. PCA plots, heatmaps, volcano plots and violin plots are generated using R packages including pcaMethods, ComplexHeatmap and ggplot2. For generating heatmaps, factoextra package was used to estimate the optimal number of clusters from k-means clustering. Gene ontology (GO) and Reactome (REAC)-based functional enrichement analysis were conducted using clusterProfiler package. Kinase activity prediction was commenced using t-statistics of phosphosites of LSS vs ST and HSS vs ST via Robust inference of Kinase Activity using functional Networks (RoKAI) webtool.

### Hydrogel constructs and encapsulation of monocytes in hydrogels

Hydrogel constructs were prepared as previously described to recapitulate the extracellular matrix composition of healthy and CAVD tissues ^30^. Hydrogels were prepared using bovine type I collagen (PureCol EZ Gel, #5074, Advanced Biomatrix) at a final concentration of 1.5 mg/mL and supplemented with chondroitin sulfate A sodium salt (#C9819, Sigma–Aldrich) dissolved in 1× PBS. The concentration of chondroitin sulfate was selected to reflect the pathological glycosaminoglycan accumulation characteristic of CAVD. Collagen hydrogels were enriched with chondroitin sulfate to a final concentration of 20 mg/mL to model severe CAVD. Following shear stress exposure, monocytes were encapsulated within the hydrogels and cultured and differentiated as described below.

### Generation of monocyte-derived macrophages

To generate monocyte-derived macrophages (MDMs), monocytes were cultured in complete RPMI-1640 medium supplemented with 50 ng/mL macrophage colony-stimulating factor (M-CSF; Miltenyi Biotec) for 2 days, with fresh medium supplied daily. To assess foam cell formation, cells were serum-starved in RPMI-1640 for 8 h on day 3 before the addition of oxidised low-density lipoprotein (oxLDL; ∼5 µg/ml, 1:200 dilution; Thermo Fisher Scientific) labelled using an Alexa Fluor 647 Protein Labelling Kit (Thermo Fisher Scientific). Cells were incubated with labelled oxLDL in serum-free medium for a further 24 h.

### PKC Activity Assay

PKC activity in monocytes was assessed using a PKC Kinase Activity Assay Kit (ab139437, Abcam) according to the manufacturer’s instructions. Briefly, monocytes were isolated as described above and lysed in extraction buffer. Protein concentration was determined using the Pierce™ BCA Protein Assay Kit (Thermo Fisher Scientific), and 10–15 µg of protein was used per assay. Samples were added to wells of the PKC substrate-coated microtiter plate, and the kinase reaction was initiated by addition of reconstituted ATP. Samples were incubated for 90 min at 30°C, after which the reaction was terminated by emptying the wells. PKC phosphospecific substrate antibody was added and incubated for 60 min at room temperature. Following washing, anti-rabbit IgG–HRP conjugate was added and incubated for 30 min at room temperature with intermittent agitation. Wells were washed before addition of TMB substrate and incubation at room temperature. The reaction was terminated by addition of stop solution, and absorbance was measured at 450 nm using a CLARIOstar Plus microplate reader (BMG LABTECH)

### Calpain activity assay

Calpain activity in monocytes was assessed using a Calpain Activity Assay Kit (ab65308, Abcam) according to the manufacturer’s instructions. Briefly, monocytes were lysed in extraction buffer, and protein concentration was determined using the Pierce™ BCA Protein Assay (Thermo Fisher Scientific). Cell lysates containing 50–200 µg of protein were adjusted to 85 µL with extraction buffer. Samples, positive controls containing active calpain, and negative controls containing calpain inhibitor were prepared in a black 96-well plate. 10 µL of 10× Reaction Buffer and 5 µL of Calpain Substrate (Ac-LLY-AFC) were added to each well, followed by incubation for 60 min at 37°C protected from light. Calpain activity was quantified by measuring fluorescence at 400 nm excitation and 505 nm emission using a CLARIOstar Plus microplate reader (BMG LABTECH).

### BrdU and MitoTracker assay

THP-1 cells were labelled with BrdU (ab142567, Abcam) prior to shear stress exposure and allowed to proliferate for 3 days. Cells were then washed with PBS and stained overnight at 4°C with a mouse anti-BrdU antibody (Cell Signaling Technology, 94079S). The following day, cells were washed and incubated with Alexa Fluor 647-conjugated anti-mouse secondary antibody (Invitrogen) for 3 h at 37°C. Cells were subsequently stained with MitoTracker Orange CMTMRos (Invitrogen, M7510), MitoTracker Green FM (Invitrogen, M7514), and Fixable Viability Dye eFluor™ 780 (Thermo Fisher Scientific, 65-0865-18). Samples were acquired using an LSRFortessa II flow cytometer (BD Biosciences), and data were analysed using FlowJo software.

### Quantitative polymerase chain reaction (qPCR)

RNAeasy Micro Kit (QIAGEN) was used to extract RNA from cells immediately after exposure to shear conditions using the manufacturer’s instructions. RNA was quantified using NanoDrop™ 2000 (Thermo Fisher Scientific). SuperScript IV VILO Master Mix (Thermo Fisher Scientific) was used to synthesize cDNA from RNA. TaqMan Fast Advanced Master Mix (Thermo Fisher Scientific) was used to detect gene expressions in concordance with the following human primers, Hs01062327_m1 *PPP1R27*, Hs01558245_m1 *TFAP4*, Hs00918966_m1 *TCF12* and Hs00167549_m1 *ANXA1* (all from Thermo Fisher Scientific). Gene expression for target genes was normalized to Hs00609296_g1 *HMBS*. For mice, Mm01241549_m1 *Piezo1* was normalized to Mm99999915_g1 *Gapdh* (both from Thermo Fisher Scientific). The fold change (2^-ΔΔCT^) was measured from at least 6 independent experiments.

### Immunocytochemistry

After exposure to shear stress and differentiation, monocytes and MDMs were fixed with 4% paraformaldehyde (Sigma-Aldrich) for 1 hr at 37°C. Then, the cells were permeabilized with 0.1% Triton X-100 (Sigma-Aldrich) for 15 mins at room temperature. Afterwards, cells were blocked with 10% normal goat serum with 5% BSA for 1 hr at 37°C and incubated in appropriate primary antibodies, 1:100 rabbit anti-CD80 (abcam-ab225674), 1:100 rabbit anti-ABCG1 (Novus Biologicals-NB400-132), 1:100 mouse anti-ABCA1 (Novus Biologicals-NB100-2068), 1:400 mouse anti-AKT (Cell Signaling Technology-#2920), and rabbit anti-phospho-AKT (Ser473) (Cell Signaling Technology-#4060) overnight at 4°C.

The following day, human cells assessed for foam cell formation and differentiation were incubated in the appropriate secondary antibodies, 1:500 anti-rabbit or anti-mouse Alexa Fluor 488, 594, or 647 (Invitrogen) or conjugated antibodies, 1:100 FITC mouse anti-human CD68 (BD Biosciences-#562117), 1:50 PE mouse anti-human CD163 (BioLegend-#556018), for 3 hrs at 37°C. For assessing AKT and phospho-AKT, 1:800 anti-mouse Alexa Fluor 594 (Invitrogen) and 1:800 anti-rabbit Alexa Fluor 647 (Invitrogen) were used. For mice, 1:100 FITC anti-mouse CD80 (#104705), 1:100 PE anti-mouse CD163 (#155307) and 1:100 APC anti-mouse F4/80 (#123115) (all from BioLegend) were used. In addition, 1:500 DAPI was used for nuclear staining, and samples were imaged using the A1R confocal microscope (Nikon).

For analysis, four fields of view were captured per each of 4-6 independent experiments. Each field of view was analyzed using NIS Elements (Nikon) by drawing regions on interest (ROIs) around at least 30 cells and the mean fluorescent intensity was recorded. The results are represented as the average value of the fields of view per independent experiment.

## Statistical analysis

Statistical analysis was conducted using GraphPad Prism 10 (GraphPad Software Inc.), and unpaired two-tailed Student’s *t*-test, or either one-way or two-way ANOVA with Tukey’s (for comparison of more than two factors), Šídák’s (for comparison of two factors) or Dunnett’s (for comparisons to the control group) post-hoc test, were used where appropriate. A p-value < 0.05 was considered significant.

## Results

### Shear stress induces global proteomic remodelling of monocytes

We previously demonstrated that pathological shear stress, similar to that experienced by circulating monocytes in patients with severe aortic stenosis, induces a pro-inflammatory phenotype through Piezo1-dependent mechanotransduction and that these effects are reversed following TAVI ^6^. To gain an unbiased understanding of the molecular signalling pathways and downstream biological processes underlying these functional responses, we investigated the effects of different shear stress magnitudes on the global monocyte proteome using quantitative data-independent acquisition (DIA) proteomics. THP-1 monocytes were exposed to physiological wall shear stress of 80 dyn/cm² (LSS) or pathological wall shear stress of 400 dyn/cm² (HSS) for 30 min before quantitative proteomic analysis (**Figure 1**). The proteomic dataset demonstrated excellent reproducibility across biological replicates, with low coefficients of variation in all experimental groups (**Figure 1a**). Principal component analysis clearly separated static, LSS and HSS samples, indicating that exposure to shear stress induced magnitude-dependent changes in the global monocyte proteome (**Figure 1b**). Hierarchical clustering further identified seven distinct protein clusters exhibiting differential protein expressions across each of these conditions (**Figure 1c**). Functional enrichment analysis demonstrated that proteins regulated by shear stress were predominantly associated with RNA processing, protein translation, mitochondrial function and inflammatory pathways (**Figure 1d**). Differential expression analysis further revealed distinct proteomic responses to low and high shear stress. Compared with static controls, exposure to LSS resulted in significant changes in proteins involved in mitochondrial translation, mitochondrial gene expression, defense responses to fungi, killing of cells of other organisms, and cytoplasmic translation. In contrast, proteins altered following HSS were primarily enriched in RNA splicing, ribonucleoprotein complex biogenesis, chromatin remodelling, translational initiation, oxidative phosphorylation, cellular respiration and ribose phosphate biosynthetic processes. Comparison of the enriched pathways between the two shear stress conditions demonstrated that while both shared alterations in translational machinery, mitochondrial function and RNA metabolism, LSS preferentially regulated pathways associated with innate immune responses and mitochondrial translation, whereas HSS predominantly affected RNA processing and translational initiation pathways. Together, these findings demonstrate that shear stress induces extensive and magnitude-dependent remodelling of the monocyte proteome.

**Figure 1.**
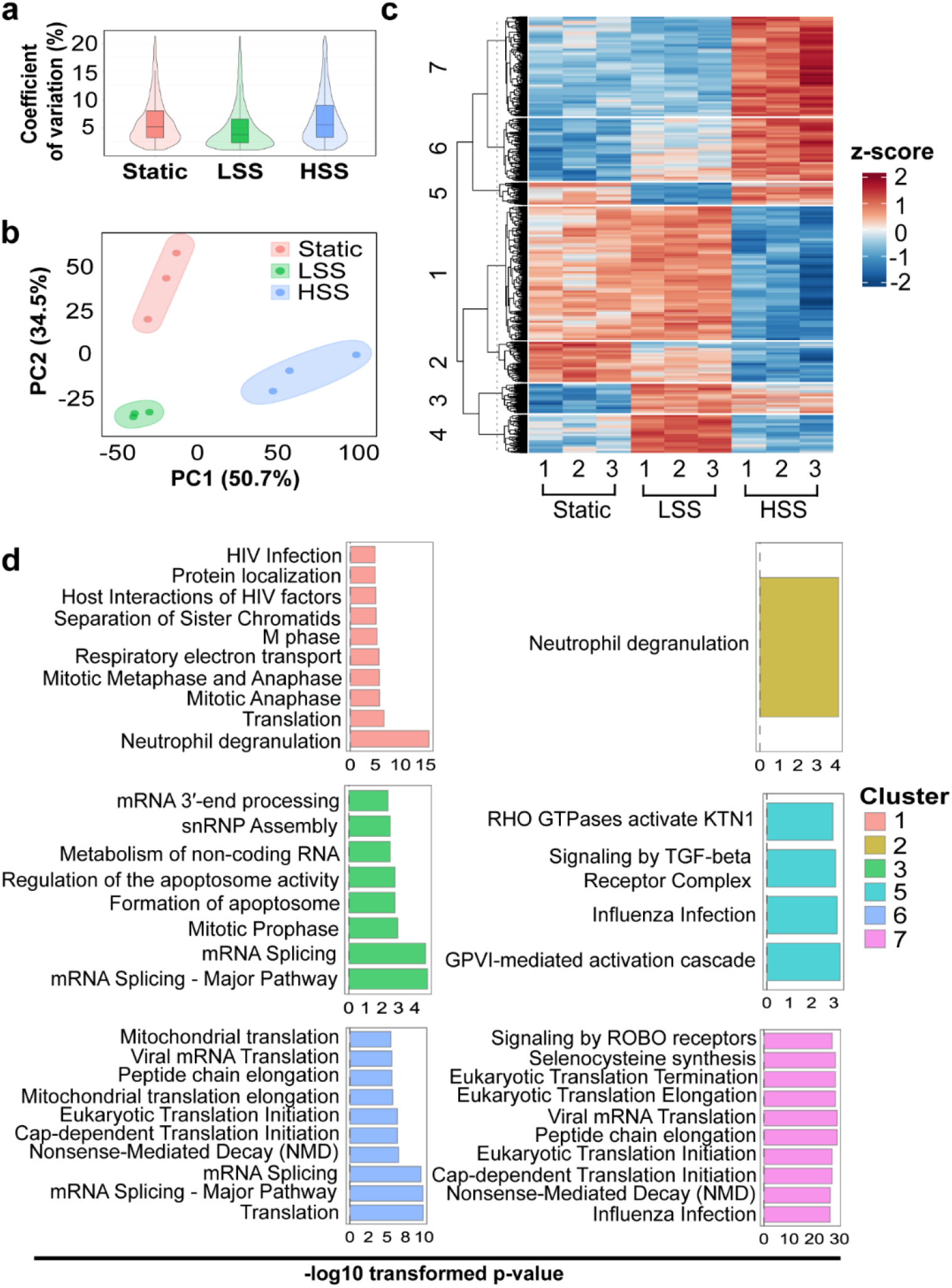
Proteomic analysis of the influence of shear stress on monocytes. **(a)** Coefficient variations of protein group Label-Free Quantification (LFQ) intensities of LSS Static, LSS and HSS groups. **(b)** Principal component analysis (PCA) of log2-transformed proteome of Static, LSS and HSS groups. **(c)** Heatmap generated based on z-score normalized protein group intensities of significantly DEPs among Static, LSS and HSS groups (One-way ANOVA, False Discovery Rate (FDR) (Benjamini–Hochberg (BH) procedure) < 0.05) with post-hoc analysis significantly different to Static. Seven signature clusters (optimal number of clusters was determined by the fviz_nbclust function through the factoextra package) were differentiated based on distance. Functional enrichment of protein groups from each cluster. **(d)** Bar graphs show the top enriched Reactome (REAC) pathways.

### Shear stress rapidly remodels the monocyte phosphoproteome

We next examined shear stress-induced changes in the phosphoproteome. More than 24,000 Class I phosphopeptides were identified across all samples, providing extensive coverage of the monocyte phosphoproteome. Similar to the global proteomic analysis, phosphoproteomic data demonstrated excellent reproducibility and clear separation of static, LSS and HSS samples by principal component analysis (**Figure 2a, b**), indicating rapid and shear-dependent remodelling of intracellular signalling pathways.

**Figure 2.**
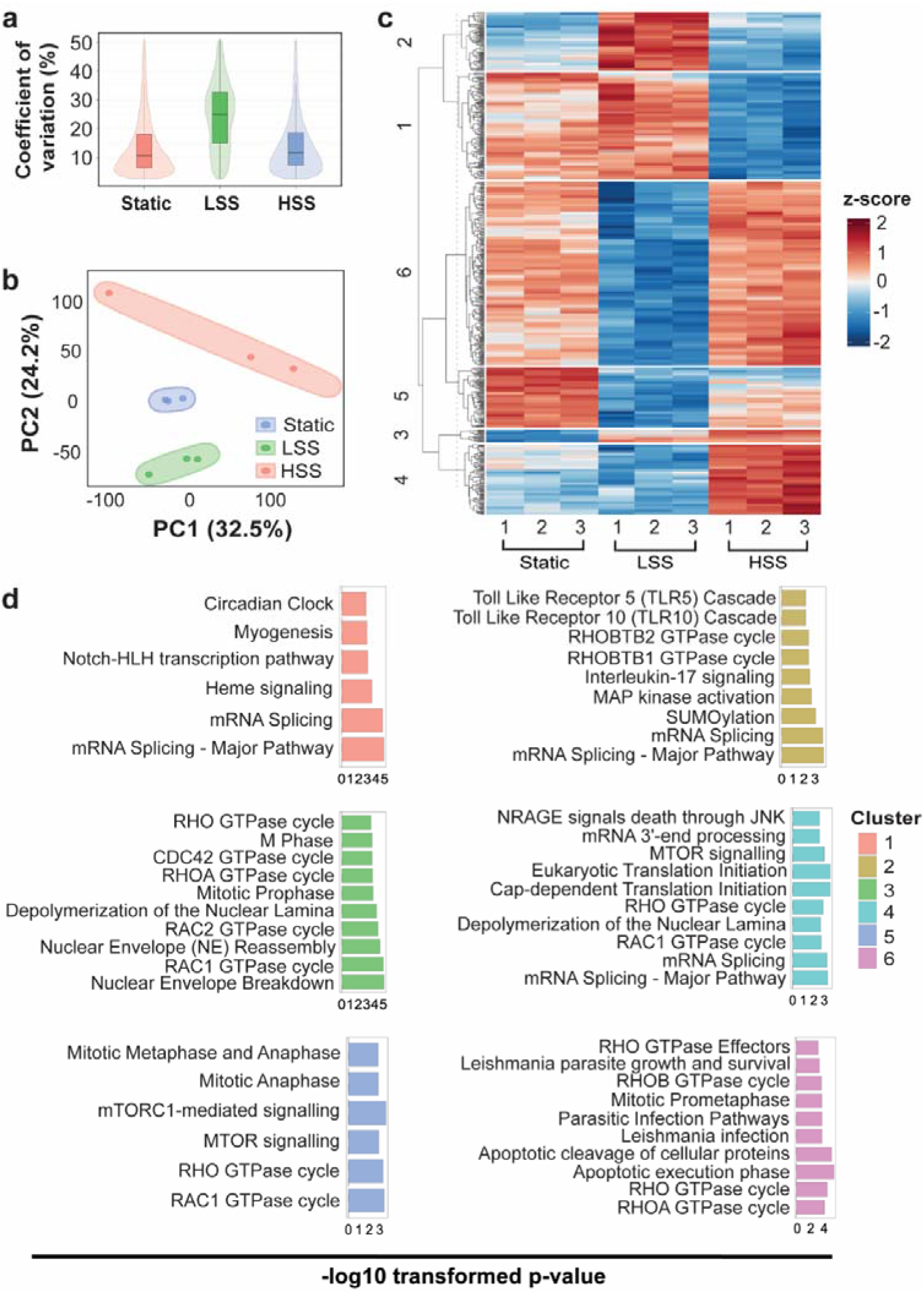
Shear stress induces distinct phosphoproteomic signatures in monocytes. **(a)** Coefficient of variation (CV) of phosphosite LFQ intensities in Static, LSS and HSS groups. **(b)** Principal component analysis (PCA) of phosphoproteomic profiles from each experimental group. **(c)** Heatmap of z-score-normalised phosphosite intensities for significantly differentially phosphorylated phosphosites across Static, LSS and HSS groups (one-way ANOVA, Benjamini–Hochberg false discovery rate (FDR) < 0.05, with post hoc comparison to Static). Six phosphorylation clusters were identified by hierarchical clustering, with the optimal number of clusters determined using the fviz_nbclust function in the factoextra R package. Functional enrichment analysis was performed for proteins corresponding to phosphosites within each cluster. The top enriched REAC pathways **(d)** are shown.

Hierarchical clustering identified seven distinct phosphorylation signatures (**Figure 2c**). Functional enrichment demonstrated that differentially phosphorylated proteins were primarily involved in RNA metabolism, mRNA splicing, translation, cytoskeletal organisation and intracellular signalling (**Figure 2d**). These findings indicate that biomechanical stimulation rapidly activates signalling pathways regulating protein synthesis, RNA metabolism and cellular organisation.

Differential phosphorylation analysis further demonstrated distinct signalling responses to different shear stress magnitudes. Compared with static controls, **LSS** significantly altered phosphosites associated with the regulation of translation, mitotic cell cycle progression, DNA damage checkpoint signalling, non-membrane-bounded organelle assembly and DNA integrity checkpoint pathways. In contrast, **HSS** predominantly regulated phosphoproteins involved in mRNA processing, nucleocytoplasmic transport, RNA localisation, chromosome organisation, chromatin remodelling, double-strand break repair and ciliary chromosome segregation. Comparison of enriched Reactome pathways between the two conditions further demonstrated that LSS preferentially modulated RHO GTPase signalling, mTORC1-mediated signalling, mitotic cell cycle, RAC1, RHOA and RHO GTPase effector pathways, whereas HSS preferentially affected mRNA splicing, RHO GTPase signalling, RAC1, RHOA, mitotic cell cycle, cellular responses to stress and apoptotic cleavage of cellular proteins. Collectively, these findings demonstrate that shear stress induces rapid, magnitude-dependent remodelling of the monocyte phosphoproteome, with differential regulation of signalling pathways governing RNA metabolism, cytoskeletal dynamics, cell cycle progression and stress-responsive signalling.

### Temporal remodelling of the monocyte proteome

Having established that different shear stress magnitudes induce distinct molecular responses, we next investigated the temporal dynamics of proteomic remodelling following exposure to pathological shear stress (400 dyn/cm²). Protein abundance remained highly reproducible across all experimental groups (**Figure 3a**), while principal component analysis demonstrated progressive separation of static, 10 min, 30 min and overnight exposure groups, indicating continuous remodelling of the monocyte proteome over time (**Figure 3b**). Hierarchical clustering identified eight distinct temporal protein expression patterns, reflecting both transient and sustained responses to shear stress (**Figure 3c**).

**Figure 3.**
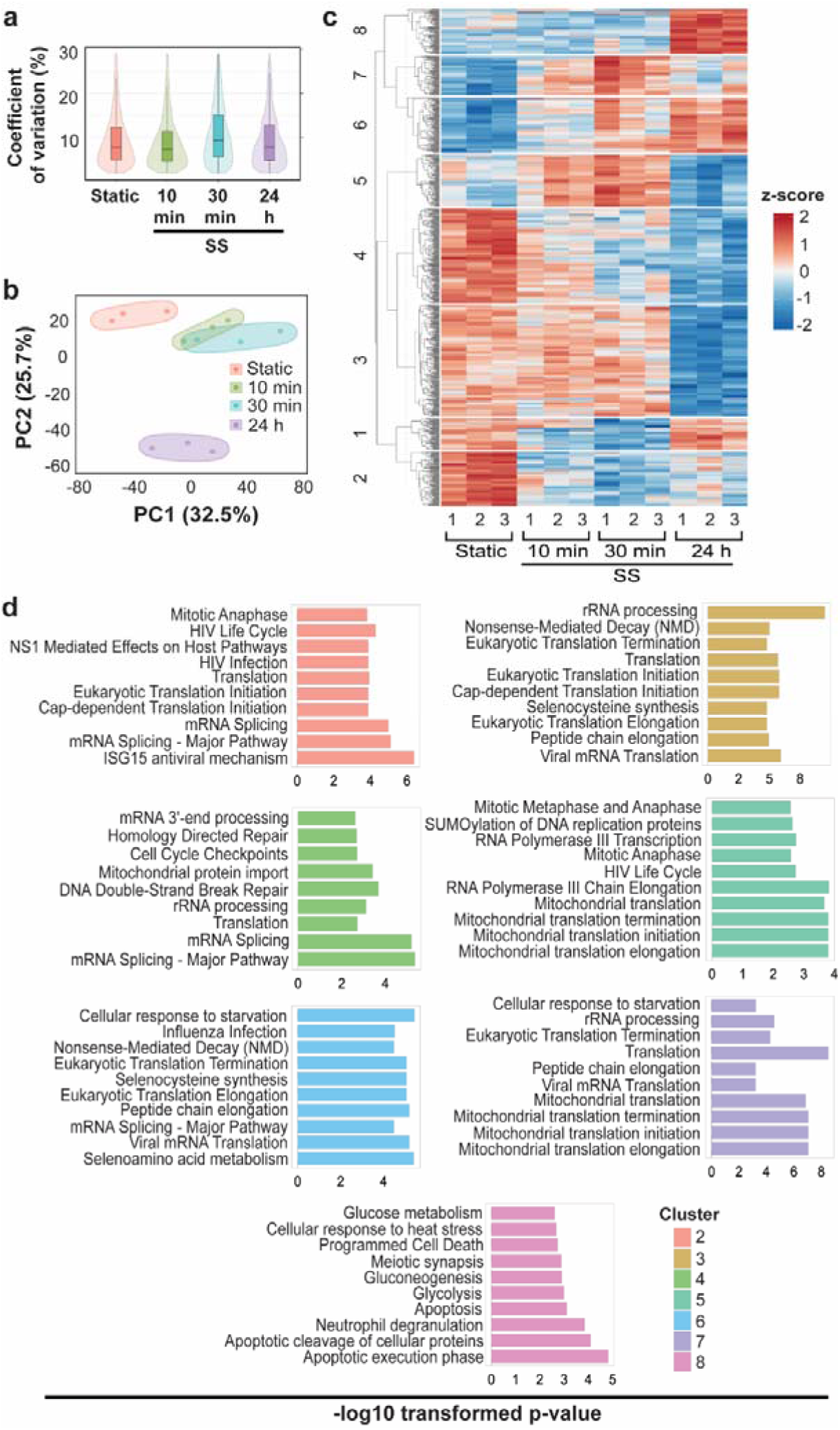
Temporal proteomic remodelling of monocytes following shear stress exposure. **(a)** Coefficient of variation (CV) of protein group LFQ intensities in Static, 10 min shear stress, 30 min shear stress and 24 h shear stress groups. **(b)** Principal component analysis (PCA) of global proteomic profiles from each experimental group. **(c)** Heatmap of z-score-normalised protein group intensities for significantly differentially expressed proteins across Static, 10 min shear stress, 30 min shear stress and ON groups (one-way ANOVA, Benjamini–Hochberg false discovery rate (FDR) < 0.05, with post hoc comparison to Static). Eight temporal protein expression clusters were identified by hierarchical clustering, with the optimal number of clusters determined using the fviz_nbclust function in the factoextra package. Functional enrichment analysis was performed for proteins within each cluster. The top enriched REAC pathways **(d)** are shown.

Functional enrichment analysis revealed that many biological processes were regulated across multiple temporal clusters, although with distinct temporal profiles (**Figure 3d**). RNA metabolism, mRNA processing, ribosome biogenesis, translation and protein targeting to the endoplasmic reticulum were among the earliest and most consistently enriched pathways, suggesting rapid reprogramming of protein synthesis immediately following shear stress exposure (Cluster 5). Mitochondrial organisation, mitochondrial gene expression, oxidative phosphorylation and aerobic respiration became increasingly enriched at later time points, indicating progressive metabolic adaptation during prolonged mechanical stimulation (Clusters 6 and 7). In parallel, pathways associated with protein targeting to mitochondria, mitochondrial translation, aerobic respiration, glucose metabolism and glycolysis were dynamically regulated throughout the time course, reflecting extensive cellular remodelling. Notably, immune-related processes, including neutrophil degranulation, activation of the innate immune system, apoptosis and glucose metabolism, were predominantly enriched following overnight exposure, suggesting that sustained shear stress promotes inflammatory and stress-response pathways after the initial adaptive response (Cluster 8). Collectively, these findings demonstrate that monocyte adaptation to shear stress occurs in a temporally coordinated manner, with an early phase dominated by translational and RNA-processing programmes followed by progressive metabolic, mitochondrial and inflammatory remodelling.

### Early phosphoregulation precedes proteomic adaptation

To investigate the kinetics of early mechanotransduction, phosphoproteomic analysis was performed on the same temporal shear stress samples used for global proteomic analysis **(Figure 4**). Phosphosite abundances were highly reproducible across all experimental groups **(Figure 4a)**, while principal component analysis demonstrated progressive separation of static, 10 min, 30 min and overnight samples, indicating continuous remodelling of the phosphoproteome over time **(Figure 4b)**. Hierarchical clustering identified seven distinct temporal phosphorylation signatures, revealing coordinated changes in phosphosite abundance throughout the course of shear stress exposure **(Figure 4c**).

**Figure 4.**
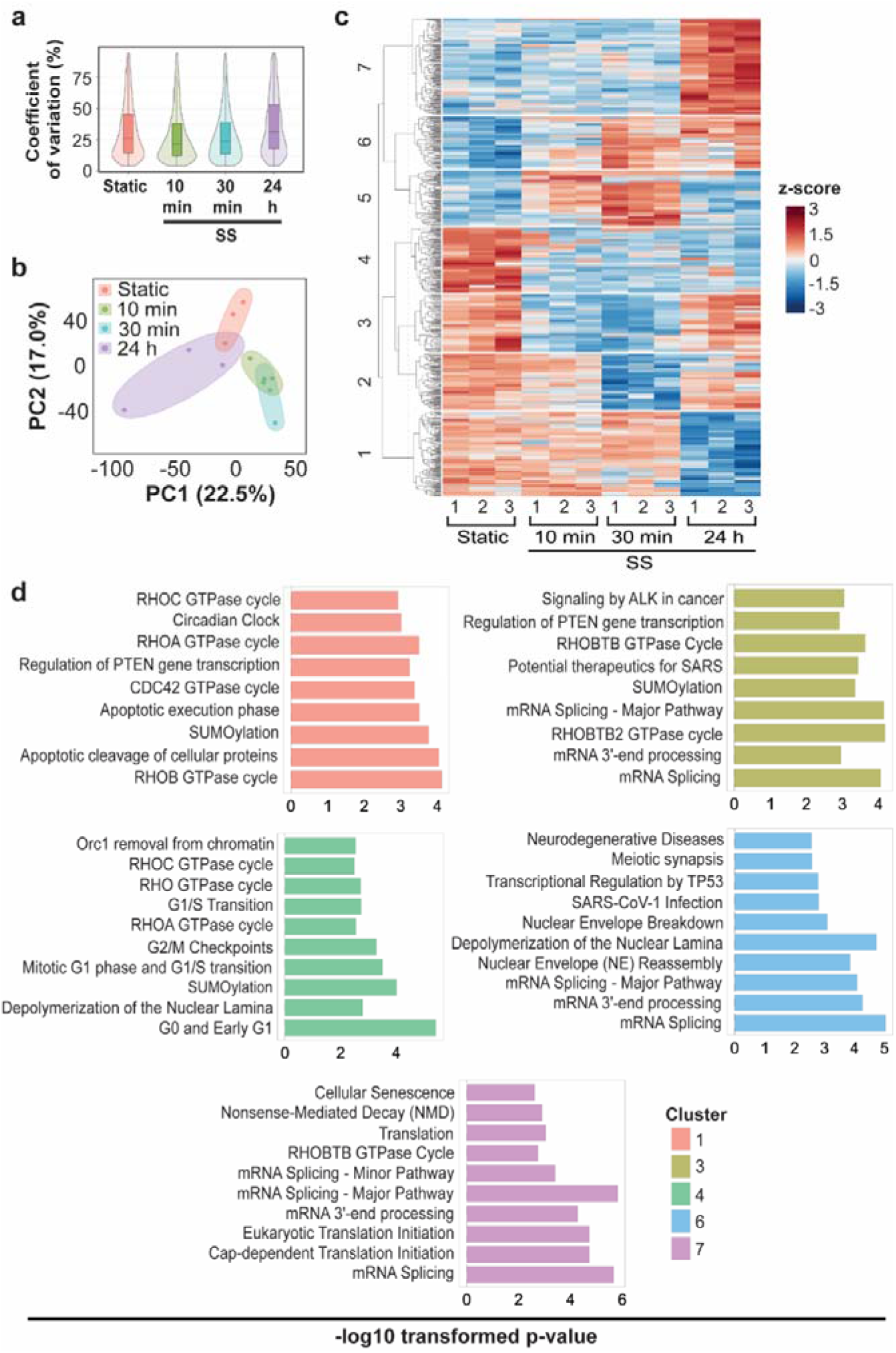
Temporal phosphoproteomic remodelling of monocytes following shear stress exposure. **(a)** Coefficient of variation (CV) of phosphosite LFQ intensities in Static, 10 min shear stress, 30 min shear stress and 24 h shear stress groups. **(b)** Principal component analysis (PCA) of phosphoproteomic profiles from each experimental group. **(c)** Heatmap of z-score-normalised phosphosite intensities for significantly differentially phosphorylated phosphosites across Static, 10 min shear stress, 30 min shear stress and 24 hours (one-way ANOVA, *P* < 0.05, with post hoc comparison to Static). Seven temporal phosphorylation clusters were identified by hierarchical clustering, with the optimal number of clusters determined using the fviz_nbclust function in the factoextra package. Functional enrichment analysis was performed for proteins corresponding to phosphosites within each cluster. The top enriched REAC pathways **(d)** are shown.

Functional enrichment analysis of the temporal phosphorylation clusters revealed dynamic regulation of multiple biological processes and signalling pathways **(Figure 4d**). Early-responsive Cluster 1 was enriched for RHO GTPase signalling, including the RHOC, RHOA, CDC42 and RHOB GTPase cycles, consistent with rapid activation of cytoskeletal remodelling and mechanotransduction pathways following shear stress (Cluster 1). Other clusters were characterised by enrichment of RNA processing, mRNA splicing, transcriptional regulation and chromatin organisation, suggesting rapid post-transcriptional and transcriptional reprogramming in response to mechanical stimulation. As shear stress exposure progressed, phosphorylation changes became increasingly associated with mRNA processing and splicing, translation initiation, nuclear organisation, TP53-mediated transcription and cellular senescence (Clusters 3, 4, 6 and 7), reflecting a transition from immediate signalling events to broader regulation of cellular adaptation. The persistent enrichment of RHO GTPase signalling, mRNA splicing, translation initiation, nuclear events mediated by TP53, cell cycle regulation and cellular senescence across multiple temporal clusters **(Figure 4d)** indicates that these signalling networks remain dynamically regulated throughout the early response to shear stress. Notably, pathways associated with RNA metabolism, translation and mitochondrial function were enriched in both the temporal phosphoproteome (**Figure 4**) and the temporal proteome (**Figure 3**), supporting the concept that early phosphorylation events initiate signalling programs that regulate change in protein abundance during monocyte adaptation to shear stress. Collectively, these findings demonstrate that phosphorylation represents an immediate and highly coordinated response to mechanical stimulation, preceding the broader changes in protein abundance observed during temporal proteomic remodelling.

### Stenosis-induced shear stress regulates actin remodeling and mitochondrial membrane potential in human monocytes

Proteomic and phosphoproteomic analyses identified shear stress-induced changes in proteins and phosphoproteins associated with cytoskeletal organisation, mitochondrial function and cellular metabolism (**Figures 1–4**). To determine whether these molecular changes were accompanied by functional alterations, we next examined the effects of physiological and stenotic shear stress on actin remodelling and mitochondrial membrane potential in human monocytes. Human monocytes isolated from the peripheral blood of healthy volunteers were exposed to LSS or HSS for 10 or 30 min and compared with static controls. We first investigated the effects of shear stress on actin cytoskeletal remodelling by quantifying the ratio of filamentous (F-actin) to globular (G-actin). Dynamic reorganisation of the actin cytoskeleton is a mechanically regulated process that governs cell morphology, deformation, migration and motility ^31^. Exposure of human monocytes to LSS for 30 min significantly increased the F-actin:G-actin ratio by 1.15 ± 0.03-fold (*P* = 0.0227) compared with static controls and by 1.20 ± 0.03-fold (*P* = 0.0054) compared with cells exposed for 10 min (**Figure 5a, b**). Similarly, exposure to HSS for 30 min increased the F-actin:G-actin ratio by 1.21 ± 0.03-fold (*P* = 0.0139) and 1.18 ± 0.03-fold (*P* = 0.0265) relative to static and 10 min, respectively (**Figure 5a, b**). These findings demonstrate that both physiological and pathological shear stress promote actin polymerisation in human monocytes, consistent with the cytoskeletal remodelling identified in the proteomic and phosphoproteomic analyses.

**Figure 5.**
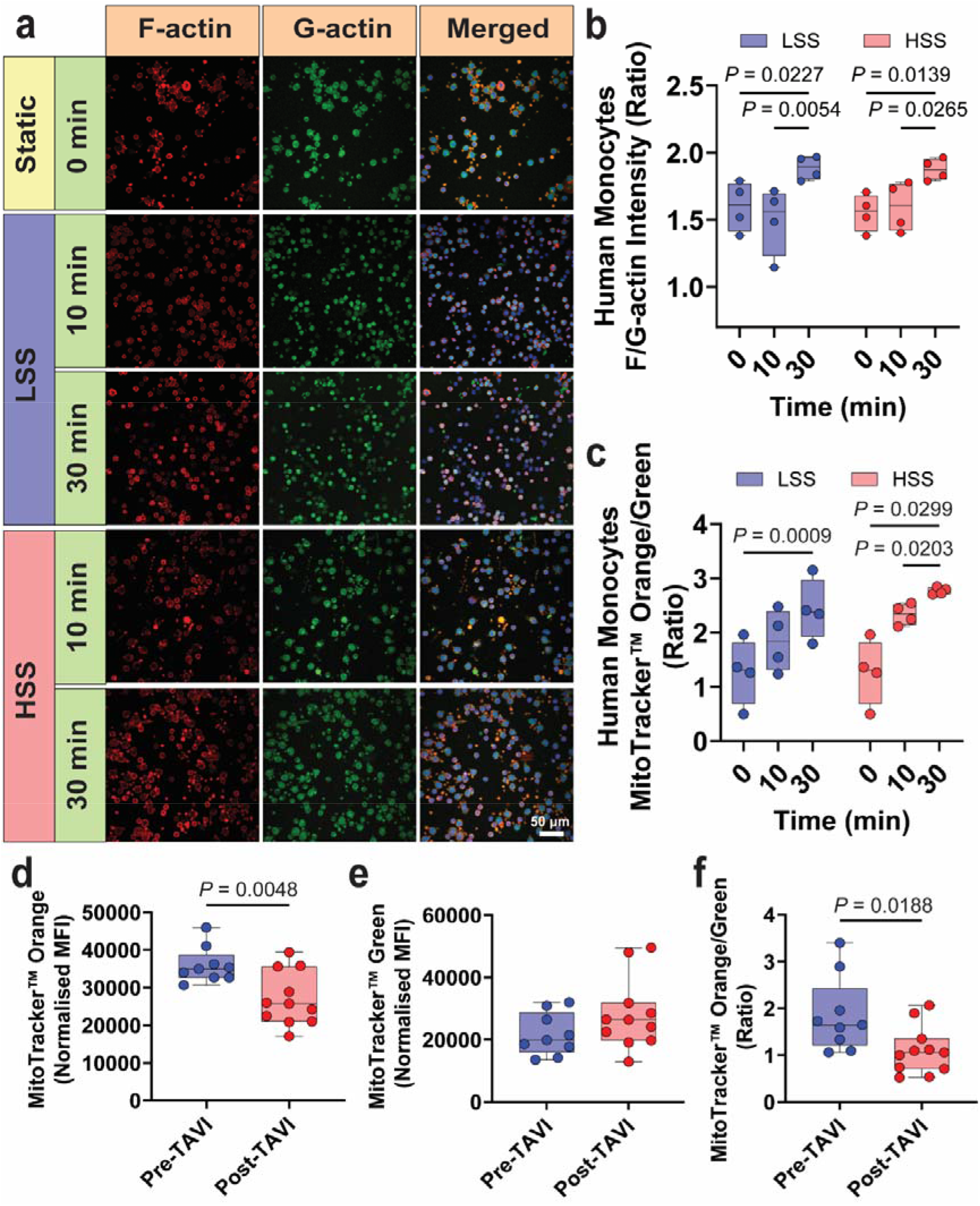
Shear stress regulates actin remodeling and mitochondrial membrane potential in human monocytes. **(a)** Representative confocal images showing F-actin (red) and G-actin (green) staining in human monocytes exposed to static conditions (0 min), low shear stress (LSS) or high shear stress (HSS) for 10 or 30 min. **(b)** Quantification of the F-actin: G-actin fluorescence intensity ratio in human monocytes under the indicated conditions (N = 4). **(c)** Quantification of the MitoTracker Orange: Green means fluorescence intensity (MFI) ratio in human monocytes exposed to the indicated conditions (N = 4). Matched two-way ANOVA with Tukey’s multiple comparisons test was used for analysis of b-c. Mean fluorescence intensity (MFI) of **(d)** MitoTracker Orange, **(e)** MitoTracker Green, and **(f)** the MitoTracker Orange: Green ratio in monocytes isolated from patients before (pre-TAVI, N = 9) and one month after transcatheter aortic valve implantation (post-TAVI, N = 11). Two-tailed unpaired t-test was used for statistical comparisons for d-f. The line in the box plot represents the median.

We next examined whether shear stress also altered mitochondrial function by assessing mitochondrial mass using MitoTracker Green and mitochondrial membrane potential using MitoTracker Orange. Following 30 min of LSS exposure, the MitoTracker Green:Orange ratio increased by 1.91 ± 0.14-fold (*P* = 0.0009) compared with static controls. A similar increase was observed following HSS exposure (2.18 ± 0.12-fold, *P* = 0.0299; **Figure 5c**), indicating that shear stress alters mitochondrial function in human monocytes.

To validate these findings in a clinically relevant setting, we took advantage of the unique opportunity provided by patients undergoing TAVI, which enabled us to compare monocytes exposed to pathological shear stress associated with severe aortic stenosis before TAVI with monocytes collected one month after TAVI, when replacement of the stenotic valve with a prosthetic valve restores near-physiological haemodynamics and markedly reduces shear stress. Following the functional responses observed in monocytes isolated from healthy volunteers, we investigated whether circulating monocytes exposed to pathological shear stress in patients with severe aortic stenosis exhibited similar mitochondrial alterations. Monocytes were isolated from patients immediately before and after TAVI, where valve replacement acutely reduces pathological shear stress from stenotic to near-physiological levels. Consistent with the *in vitro* observations, MitoTracker Orange fluorescence was 1.33 ± 0.08-fold higher (*P* = 0.0048) in monocytes collected before TAVI than after TAVI, while the MitoTracker Orange: Green ratio was 1.69 ± 0.16-fold higher (*P* = 0.0188) before TAVI (**Figure 5d–f**). These findings confirm in a clinical cohort that exposure to stenosis-associated shear stress is associated with increased mitochondrial membrane potential, supporting the physiological relevance of the *in vitro* observations.

### Pathological shear stress increases PKC-family activity and reduces AKT Ser473 phosphorylation in monocytes

To identify the signalling pathways driving the phosphoproteomic remodelling induced by pathological shear stress, we performed kinase activity prediction analysis using significantly regulated phosphosites. Kinase activity prediction revealed extensive remodelling of kinase signalling following HSS exposure, with several kinases predicted to be either activated or inhibited **(Figure 6a)**. Among these, PKCι was predicted to have increased activity, whereas AKT1, AKT2 and AKT3 were predicted to have reduced activity, identifying PKCι and AKT signalling as candidate responses to HSS. In contrast, LSS was associated with reduced predicted activity of multiple PKC isoforms, including PKCα, PKCβII, PKCδ, PKCε, PKCγ, PKCι, PKCθ and PKCζ, together with AKT1, AKT2 and AKT3. To investigate signalling responses associated with the kinase predictions, we measured intracellular calcium, AKT Ser473 phosphorylation and total PKC activity. Intracellular calcium was assessed because it can regulate calpain and other calcium-responsive signalling pathways. HSS for 30 min significantly increased intracellular calcium levels by 1.44 ± 0.06-fold compared with static controls (*P* = 0.0049) **(Figure 6b, c)**. Following 30 min of HSS, the ratio of AKT phosphorylated at Ser473 to total AKT was reduced by approximately 10% compared with static controls (*P* = 0.0178; **Figure 6b, d**). This finding is consistent with the predicted reduction in AKT activity, although phosphorylation at a single site does not establish the activity of individual AKT isoforms.

**Figure 6.**
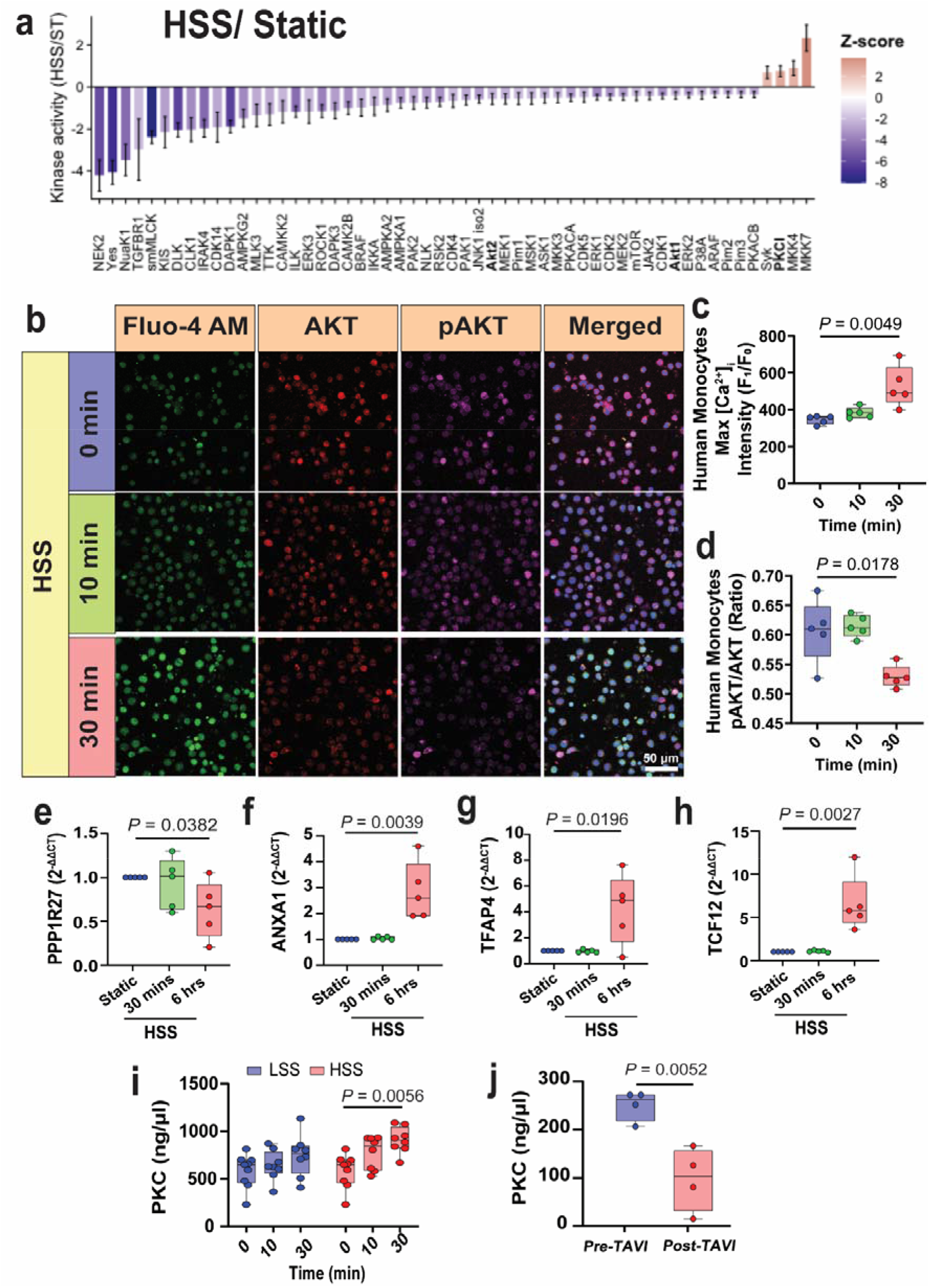
Shear stress alters calcium, PKC-family activity and AKT phosphorylation in monocytes. **(a)** Predicted upstream kinase activity in THP-1 cells exposed to HSS compared with static (ST) conditions, determined using the RoKAI web tool. **(b)** Representative confocal images of intracellular calcium ([Ca²]i; Fluo-4 AM, green), total AKT (red) and phosphorylated AKT (Ser473) (pAKT, magenta) in human monocytes exposed to HSS for 0 (static), 10 or 30 min. Quantification of **(c)** intracellular calcium ([Ca²]i) levels and **(d)** the pAKT/AKT ratio (N = 5). **(e–h)** Relative gene expression of PPP1R27, ANXA1, TFAP4, and TCF12 in THP-1 cells under static conditions or following exposure to HSS for 30 min or 6 h (N = 5). The data in b-h were analyzed using matched one-way ANOVA with Dunnett’s multiple comparisons test. **(i)** PKC activity in primary human monocytes maintained under static conditions (0 min, N = 9) or exposed to LSS or HSS for 10 (N = 8) or 30 min (N = 8). Data was analyzed using unmatched two-way ANOVA with Tukey’s multiple comparisons test. **(j)** PKC activity in circulating monocytes isolated from patients before (pre-TAVI, N = 4) and one month after transcatheter aortic valve implantation (post-TAVI, N = 4). Data was analyzed using two-tailed unpaired t-test. The line in the box plots represents the median.

We next measured total PKC-family activity in primary human monocytes. HSS for 30 min increased PKC activity by 1.55 ± 0.09-fold compared with static controls (*P* = 0.0056; **Figure 6i**) although activity did not differ significantly between HSS and LSS. Together, these measurements show increased intracellular calcium and PKC-family activity, alongside a modest reduction in AKT Ser473 phosphorylation, following HSS exposure.

We next investigated whether the kinase pathways identified by phosphoproteomic analysis contributed to the functional responses of monocytes to HSS. As the phosphoproteomic analysis was performed in THP-1 cells, kinase inhibition studies were initially conducted in the same cell model. HSS markedly reduced THP-1 cell proliferation, and pharmacological inhibition of PKC or AKT modified this response, supporting a functional role for these pathways in the cellular response to pathological shear stress. PKC and AKT inhibition also differentially altered mitochondrial responses to HSS, with AKT inhibition predominantly affecting mitochondrial membrane potential and PKC inhibition affecting mitochondrial mass and the membrane potential-to-mass ratio.

We additionally examined cyclin-dependent kinases and casein kinase 2, which were implicated by the phosphoproteomic analysis. Inhibition of these kinases attenuated the HSS-induced reduction in proliferation and altered mitochondrial membrane potential, without significantly affecting mitochondrial mass or the membrane potential-to-mass ratio. Together, these pharmacological studies support a functional contribution of the kinase networks identified by phosphoproteomics to the proliferative and mitochondrial responses of monocytes to HSS.

As these signalling pathways can regulate downstream transcriptional responses, we next examined the mRNA expression of selected genes identified from the phosphoproteomic analysis. HSS significantly decreased PPP1R27 mRNA expression at 6 h compared with static (*P* = 0.0382; **Figure 6e**). ANXA1 mRNA expression increased 2.85 ± 0.41-fold following 6 h of HSS (*P* = 0.0039), whereas no significant change was observed after 30 min (**Figure 6f**). Similarly, the mRNA expression of the transcription factors TFAP4 and TCF12 was significantly increased following 6 h of HSS (4.24 ± 0.98-fold, *P* = 0.0198 and 6.55 ± 1.17-fold, *P* = 0.00027, respectively; **Figure 6g, h**). These findings indicate that early phosphorylation changes induced by HSS are followed by changes in the expression of selected downstream genes. Finally, to determine whether these signalling changes were reflected in patients with severe aortic stenosis, we measured PKC activity in circulating monocytes isolated before and one month after TAVI. PKC activity was decreased to 0.39 ± 0.14-fold of pre-TAVI levels (*P* = 0.0052) **(Figure 6j)**. Although these responses differed from those observed following acute HSS stimulation *in vitro*, they likely reflect long-term haemodynamic adaptation after relief of pathological shear stress rather than the acute mechanotransduction responses captured in the in vitro model. Collectively, these findings demonstrate that pathological shear stress activates a calcium-dependent mechanotransduction pathway characterised by increased intracellular calcium, activation of PKC and calpain, and suppression of AKT signalling. These early signalling events are accompanied by subsequent transcriptional changes in mechanosensitive genes and highlight distinct adaptations occurring during chronic haemodynamic remodelling in patients with aortic stenosis.

### Shear stress and a CS-enriched CAVD-like extracellular matrix regulate oxLDL uptake and macrophage phenotype

To investigate whether the mechanical history of circulating monocytes influences their differentiation following recruitment into the aortic valve, we utilised our previously established three-dimensional CS-enriched CAVD-like extracellular matrix model ^30^. This model incorporates collagen hydrogels supplemented with chondroitin sulfate (CS) to mimic the glycosaminoglycan-rich extracellular matrix characteristic of CAVD **(Figure 7a).** Human monocytes were exposed to pathological HSS for 30 min or 6 h before being embedded in collagen hydrogels in the presence or absence of CS. Cells were subsequently differentiated into macrophages over 2 days prior to downstream analyses. We then assessed foam cell formation by measuring oxidised low-density lipoprotein (oxLDL) uptake together with the cholesterol efflux transporters ATP-binding cassette transporter G1 (ABCG1) and ATP-binding cassette transporter A1 (ABCA1).

**Figure 7.**
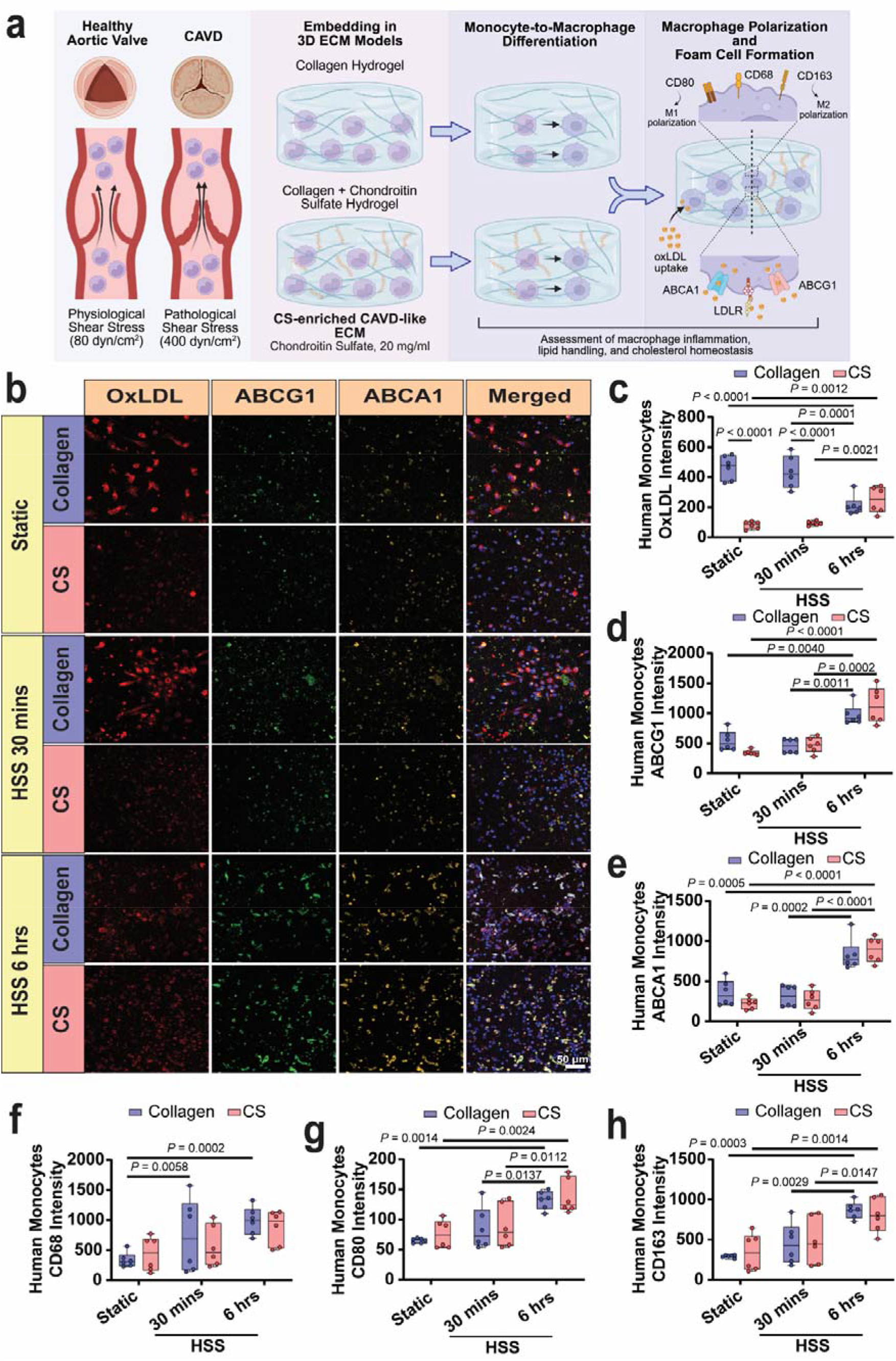
Pathological shear stress and ECM remodelling regulate oxLDL uptake, macrophage differentiation, and polarization-associated marker expression. (a) Schematic representation of experimental approach. (b) Representative images of oxLDL (red), ABCG1 (green), and ABCA1 (yellow) in human monocytes exposed to HSS and cultured into ECM model of CAVD (collagen hydrogels with or without 20µg/ml of chondroitin sulfate (CS). Graphs showing (c) oxLDL, (d) ABCG1, (e) ABCA1, (f) CD68, (g) CD80, (h) CD163) fluorescent intensities in human monocytes exposed to the above conditions (N = 6). Data was analyzed using matched two-way ANOVA with Tukey’s multiple comparisons test. The line in the box plots represent the median.

HSS exposure for 30 min had no significant effect on oxLDL uptake compared with static controls (**Figure 7b, c**). In contrast, monocytes exposed to HSS for 6 h before differentiation exhibited a 0.41 ± 0.06-fold reduction in oxLDL uptake compared with static controls (*P* < 0.0001) and a 0.45 ± 0.06-fold reduction compared with HSS for 30 min (*P* = 0.0001) when cultured in collagen hydrogels alone (**Figure 7b, c**).

Incorporation of CS into the collagen hydrogels further altered macrophage lipid uptake. Under static conditions, CS reduced oxLDL uptake by 0.18 ± 0.07-fold compared with collagen alone (*P* < 0.0001). Similarly, macrophages differentiated from monocytes exposed to HSS for 30 min exhibited a 0.22 ± 0.06-fold reduction in oxLDL uptake in the presence of CS (*P* < 0.0001) (**Figure 7b, c**). In contrast, following 6 h of HSS, CS no longer reduced oxLDL uptake relative to collagen alone. Instead, oxLDL uptake was 2.30 ± 0.16-fold higher than static monocytes (*P* = 0.0012) and 2.08 ± 0.14-fold higher than monocytes exposed to HSS for 30 min (*P* = 0.0021) (**Figure 7b, c**), suggesting that prolonged exposure to pathological shear stress alters the macrophage response to a CS-enriched extracellular matrix.

To determine whether these changes in lipid uptake were accompanied by alterations in cholesterol efflux pathways, we next examined expression of ABCG1 and ABCA1. In collagen hydrogels without CS, HSS for 6 h increased ABCG1 expression by 1.42 ± 0.09-fold compared with static controls (*P* = 0.0040) and by 1.93 ± 0.09-fold compared with HSS for 30 min (*P* = 0.0011) (**Figure 7b, d**). Similarly, in the presence of CS, ABCG1 expression increased by 2.24 ± 0.08-fold (*P* < 0.0001) and 1.76 ± 0.08-fold (*P* = 0.0002) compared with static and HSS for 30 min, respectively (**Figure 7b, d**). A similar pattern was observed for ABCA1. HSS for 6 h increased ABCA1 expression by 1.89 ± 0.09-fold (*P* = 0.0005) and 2.52 ± 0.09-fold (*P* = 0.0002) compared with static and HSS for 30 min, respectively (**Figure 7b, e**). In the presence of CS, ABCA1 expression increased by 2.80 ± 0.10-fold and 2.36 ± 0.10-fold compared with static and HSS for 30 min, respectively (both *P* < 0.0001) (**Figure 7b, e**). However, supplementation of the collagen hydrogels with CS alone did not significantly alter ABCG1 or ABCA1 expression compared with collagen-only controls (**Figure 7b, d-e**).

Next, we assessed expression of macrophage differentiation and polarization-associated markers. CD68 was used as a pan-macrophage marker, whereas CD80 and CD163 were assessed as markers associated with pro-inflammatory and anti-inflammatory macrophage phenotypes, respectively. Monocytes exposed to HSS for 6 h and differentiated within collagen hydrogels exhibited a marked increase in CD68 expression, with levels increasing by 3.34 ± 0.14-fold compared with static controls (*P* = 0.0002) and by 2.73 ± 0.16-fold compared with HSS for 30 min (*P* = 0.0058) **(Figure 7f)**. These increases were not observed when macrophages were differentiated within the CS-enriched extracellular matrix **(Figure 7f,)**. Similarly, HSS for 6 h increased CD80 expression by 2.10 ± 0.09-fold compared with static controls (*P* = 0.0014) and by 1.63 ± 0.09-fold compared with HSS for 30 min (*P* = 0.0137) in collagen hydrogels **(Figure 7g)**. Within the CS-enriched matrix, HSS for 6 h also increased CD80 expression by 1.38 ± 0.08-fold compared with static controls (*P* = 0.0024) and by 1.49 ± 0.08-fold compared with HSS for 30 min (*P* = 0.0112) **(Figure 7g)**. Expression of CD163 followed a similar pattern. In collagen hydrogels, HSS for 6 h increased CD163 expression by 2.96 ± 0.11-fold compared with static controls (*P* = 0.0003) and by 1.85 ± 0.11-fold compared with HSS for 30 min (*P* = 0.0029) **(Figure 7h)**. Likewise, within the CS-enriched matrix, HSS for 6 h increased CD163 expression by 1.46 ± 0.10-fold compared with static controls (*P* = 0.0014) and by 1.59 ± 0.10-fold compared with HSS for 30 min (*P* = 0.0147) **(Figure 7h)**. Collectively, these findings demonstrate that prolonged exposure to pathological shear stress conditions circulating monocytes to alter their subsequent macrophage phenotype within a CAVD-like extracellular environment. Pathological shear stress modulated macrophage lipid handling and foam cell formation and altered the expression of differentiation-and polarization-associated markers, whereas incorporation of chondroitin sulfate into the extracellular matrix had comparatively little additional effect on macrophage marker expression. The concurrent increase in both CD80 and CD163 suggests that pathological shear stress promotes a complex macrophage activation state rather than a simple shift towards a single polarization phenotype. Together, these findings suggest that the haemodynamic history of circulating monocytes may influence their functional phenotype following recruitment into the diseased aortic valve.

### PKC signalling negatively regulates oxLDL uptake and modulates macrophage differentiation following pathological shear stress

Our phosphoproteomic analysis and upstream kinase prediction identified PKC signalling as one of the pathways activated in monocytes following exposure to pathological shear stress (**Figure 6**). To determine whether this signalling pathway contributes to the altered macrophage phenotype induced by shear conditioning, we next investigated the functional role of PKC in regulating lipid uptake. Human monocytes were exposed to HSS for 6 h and differentiated into macrophages in collagen hydrogels in the presence or absence of the broad-spectrum PKC inhibitor Calphostin C or Rottlerin, an inhibitor with greater selectivity towards the PKCδ isoform ^32^.

In collagen hydrogels, inhibition of PKC with Calphostin C significantly increased oxLDL uptake by 1.58 ± 0.10-fold compared with vehicle-treated controls (*P* = 0.0159) (**Figure 8a, b**). Similarly, inhibition of PKCδ with Rottlerin increased oxLDL uptake by 1.59 ± 0.10-fold (*P* = 0.0283) **(Figure 8a, b)**. In contrast, neither inhibitor significantly altered oxLDL uptake when macrophages were differentiated within the CS-enriched extracellular matrix **(Figure 8a, b)**. To determine whether PKC also regulates cholesterol efflux pathways, we next examined the expression of the cholesterol transporters ABCG1 and ABCA1. Broad PKC inhibition with Calphostin C significantly increased ABCG1 expression by 1.40 ± 0.08-fold in collagen hydrogels (*P* = 0.0039) and by 1.46 ± 0.07-fold in the CS-enriched matrix (*P* = 0.0016) **(Figure 8a, c)**. Likewise, Calphostin C increased ABCA1 expression by 1.96 ± 0.07-fold (*P* < 0.0001) in collagen hydrogels and 1.55 ± 0.09-fold (*P* = 0.0020) in the presence of CS **(Figure 8a, d)**. In contrast, inhibition of PKCδ with Rottlerin did not significantly alter the expression of either ABCG1 or ABCA1 under either extracellular matrix condition **(Figure 8a, d)**.

**Figure 8.**
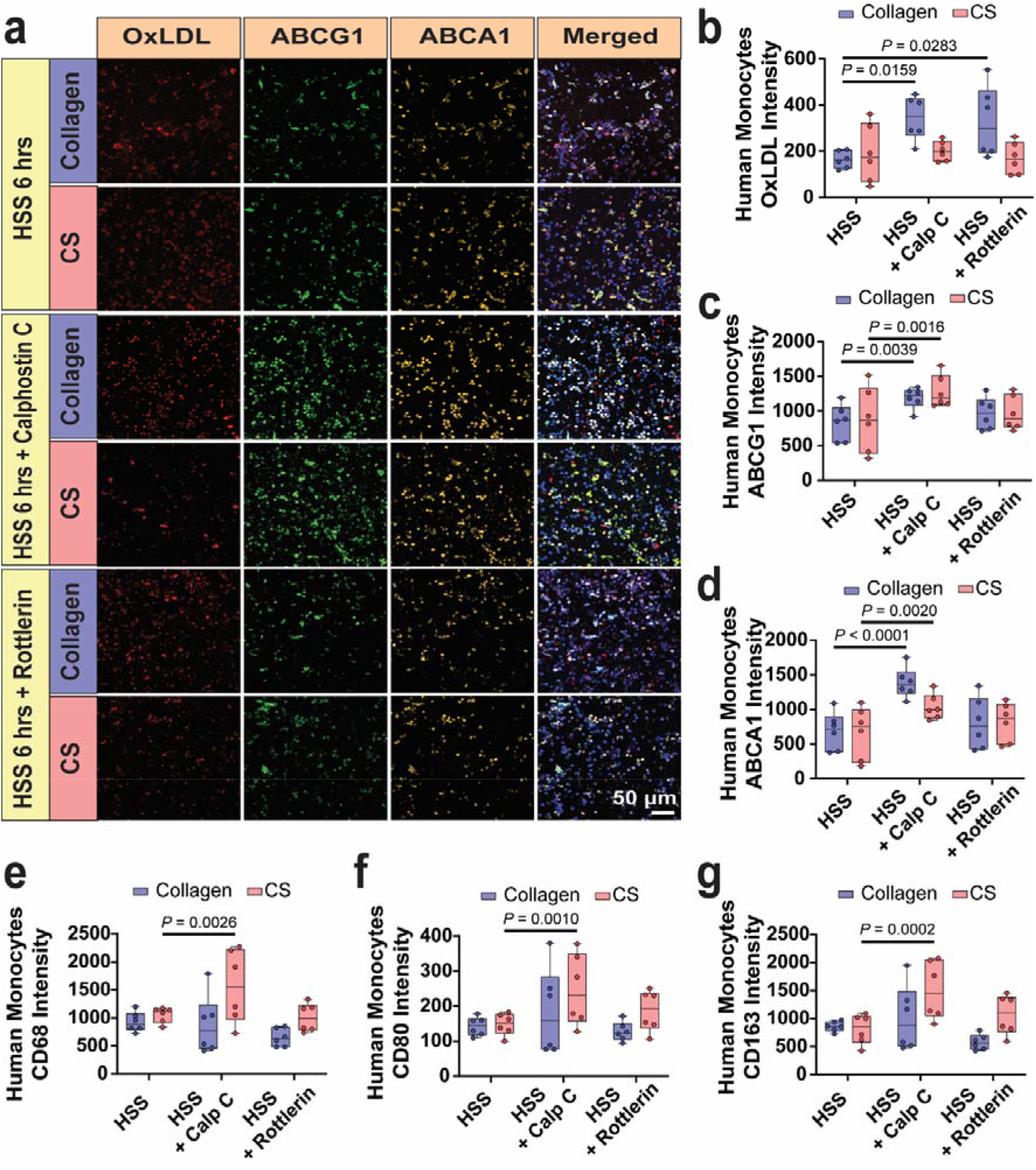
Inhibition of PKCs increases oxLDL uptake in monocyte-derived macrophages exposed to HSS. **(a)** Representative images of oxLDL (red), ABCG1 (green), and ABCA1 (yellow) in human monocytes exposed to HSS and cultured on collagen hydrogels with or without chondroitin sulfate (CS), and addition of 10 nM Calphostin C (Calp C) or 2 µM Rottlerin. Graphs showing **(b)** oxLDL uptake, **(c)** ABCG1, **(d)** ABCA1, **(e)** CD68, **(f)** CD80, and **(g)** CD163 fluorescent intensities in human monocytes exposed to the above conditions (N = 6). Data was analyzed using matched two-way ANOVA with Tukey’s multiple comparisons test. The line in the box plots represent the median.

Next, we investigated whether PKC also contributes to macrophage differentiation induced by shear-conditioned monocytes. Human monocytes exposed to HSS for 6 h were differentiated into macrophages in collagen hydrogels in the presence or absence of the broad-spectrum PKC inhibitor Calphostin C or the PKCδ-selective inhibitor Rottlerin, and expression of macrophage differentiation and polarization-associated markers was assessed. In collagen hydrogels, neither Calphostin C nor Rottlerin significantly altered the expression of the pan-macrophage marker CD68 or the polarization-associated markers CD80 and CD163 compared with vehicle-treated controls (**Figure 8e-g**). In contrast, within the CS-enriched extracellular matrix, broad PKC inhibition with Calphostin C significantly increased CD68 expression by 1.48 ± 0.10-fold (*P* = 0.0026), CD80 expression by 1.48 ± 0.11-fold (*P* = 0.0010), and CD163 expression by 1.65 ± 0.09-fold (*P* = 0.0002) compared with control macrophages. Inhibition of the PKCδ isoform with Rottlerin did not significantly affect the expression of these markers under either extracellular matrix condition (**Figure 8e-g**). Collectively, these findings demonstrate that PKC signalling functions downstream of pathological shear stress to regulate both macrophage lipid handling and differentiation within a CS-enriched CAVD-like extracellular matrix.

### Shear stress response of mouse monocytes are dependent on Piezo1-induced calcium transients and calpain

We next investigated whether the calpain and PKC signalling responses to pathological shear stress are mediated by the mechanosensitive ion channel Piezo1. Given the established role of Piezo1 in monocyte mechanotransduction under pathological shear stress ^5^, we hypothesised that Piezo1 acts upstream of the signalling pathways identified in the present study. To investigate its role, we generated a myeloid-specific Piezo1 knockout mouse model (*Piezo1*^Δ*LysM*^) by crossing *Piezo1^flox/flox^* mice with LysM-Cre mice, enabling selective deletion of Piezo1 in myeloid cells. Bone marrow-derived monocytes were isolated from *Piezo1^flox/flox^*wild-type (WT) littermates and *Piezo1*^Δ*LysM*^ knockout (KO) mice for all subsequent experiments. The genotype of each mouse was confirmed by sequencing of tail DNA, followed by validation of Piezo1 deletion at both the transcript and functional levels.

Successful deletion of Piezo1 was confirmed by qPCR, which demonstrated a 0.48 ± 0.06-fold reduction in Piezo1 mRNA expression in KO monocytes compared with WT controls (*P* < 0.0001) (**Figure 9a**). Functional validation of Piezo1 deletion was obtained by measuring intracellular calcium responses following stimulation with the Piezo1 agonist Yoda1 (50 μM). Consistent with successful disruption of Piezo1 function, KO monocytes exhibited a 0.49 ± 0.08-fold reduction in intracellular calcium transients compared with WT monocytes (*P* = 0.0009) (**Figure 9b, c**).

**Figure 9.**
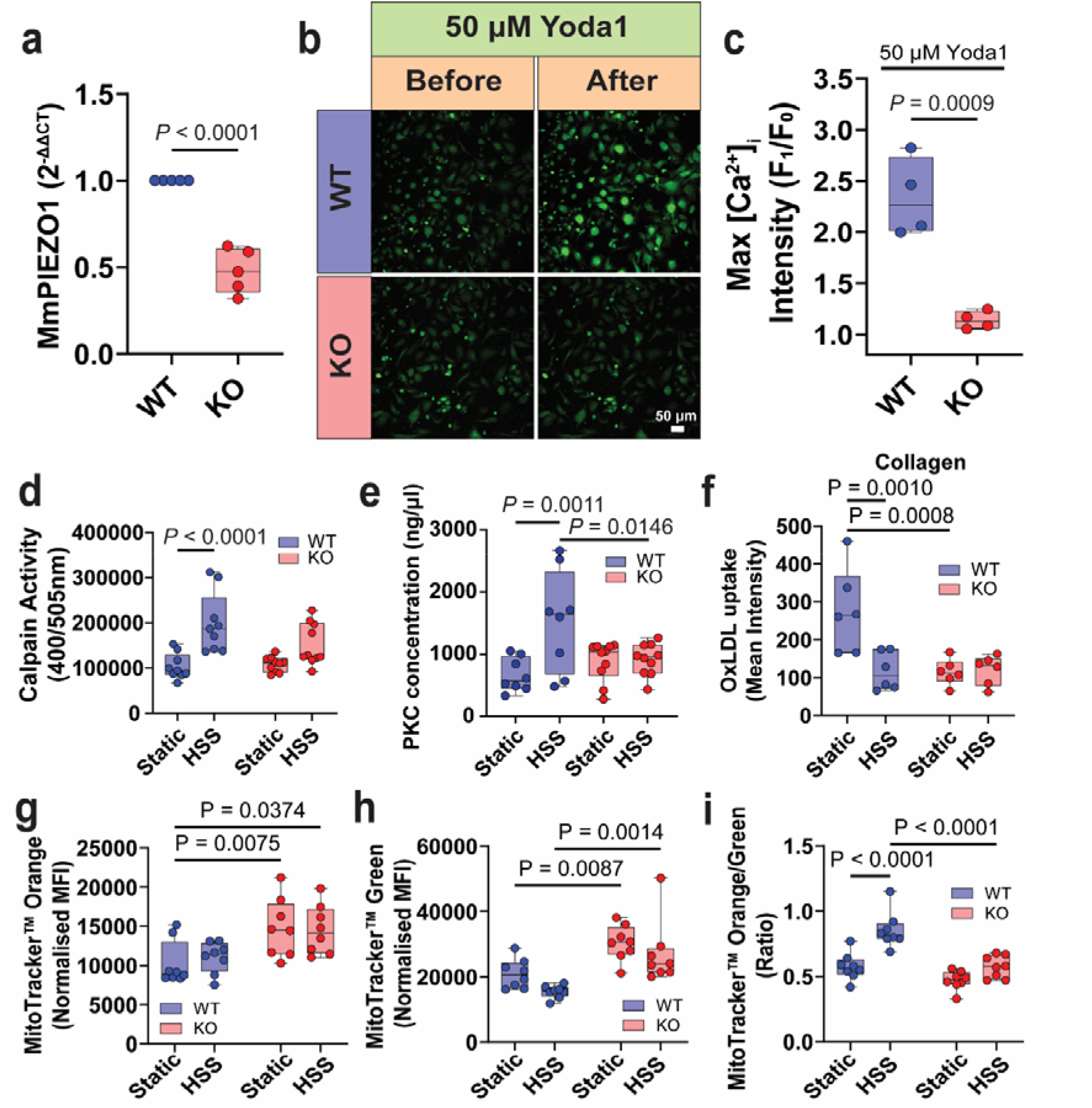
Piezo1 mediates stress-induced calcium signalling and downstream calpain activation in bone marrow-derived monocytes. **(a)** Piezo1 mRNA expression in bone marrow-derived monocytes isolated from *Piezo1^flox/flox^* wild-type (WT) and *Piezo1*^Δ*LysM*^ knockout (KO) mice (N = 5). Data was analyzed using two-tailed unpaired t-test. **(b)** Representative Fluo-4 AM calcium imaging of WT and Piezo1-KO monocytes following stimulation with the Piezo1 agonist Yoda1 (50 μM). **(c)** Quantification of the maximum intracellular calcium response ([Ca²□]□) following Yoda1 stimulation (N = 4). Data was analyzed using two-tailed unpaired t-test. **(d)** Calpain activity (WT, N = 9 and KO, N = 10) and **(e)** PKC activity (WT, N = 8 and KO, N = 10) in WT and Piezo1-KO monocytes under static conditions or following exposure to high shear stress (HSS). Quantification of **(f)** MitoTracker Orange fluorescence, **(g)** MitoTracker Green fluorescence, and **(h)** the ratio of mitochondrial membrane potential to mitochondrial mass (MitoTracker Orange/Green) in WT and Piezo1-KO monocytes under static conditions or following HSS exposure (N = 8). Data in d-i was analyzed using unmatched two-way ANOVA with Šídák’s multiple comparisons test. The line in the box plots represent the median.

We next investigated whether Piezo1 regulates the downstream signalling pathways identified by our phosphoproteomic analyses. Consistent with the predicted activation of PKC signalling, HSS-induced PKC activation was dependent on Piezo1. HSS exposure markedly increased PKC activity in WT monocytes compared to Piezo1-deficient monocytes (*P* = 0.0146) (**Figure 9e**), indicating that Piezo1 is required for normal PKC activation in response to mechanical stimulation and that loss of Piezo1 alters basal PKC signalling dynamics.

Additionally, we measured foam cell formation from monocyte-derived macrophages cultured in collagen hydrogels in which HSS reduced oxLDL uptake in WT monocytes compared to static (*P* = 0.0010), and Piezo1 KO attenuated oxLDL uptake in static conditions (*P* = 0.0008) (**Figure 9f**), suggesting that Piezo1 may mediate foam cell differentiation.

Because mitochondrial pathways were among the most significantly enriched biological processes identified by the proteomic analyse between static and both of shear stress groups, we next examined whether Piezo1 contributes to mitochondrial adaptations following pathological shear stress. Piezo1 deletion resulted in increased mitochondrial membrane potential (MitoTracker Orange) in static (*P* = 0.0075) and HSS (*P* = 0.0374), in addition to mitochondrial mass (MitoTracker Green) in static (*P* = 0.0087) and HSS (*P* = 0.0014) (**Figure 9g–i**), and the ratio of mitochondrial membrane potential to mitochondrial mass was significantly increased in WT monocytes by HSS (*P* < 0.0001), and reduced following HSS in Piezo1-deficient monocytes compared with WT monocytes (*P* < 0.0001), indicating that Piezo1 contributes to mitochondrial responses to mechanical stimulation.

Finally, given that calpain is a well-established downstream effector of Piezo1-mediated calcium influx, we assessed whether shear stress-induced calpain activation was also dependent on Piezo1. In monocytes isolated from wild-type mice, exposure to HSS significantly increased calpain activity. In contrast, Piezo1-deficient monocytes showed no significant increase in calpain activity following HSS (**Figure 9d**).

## Discussion

CAVD is characterised by progressive valvular fibrosis and calcification, ultimately leading to altered haemodynamics and severe aortic stenosis ^33, 34^. Although inflammation is recognized as a major contributor to disease progression ^35^, it remains unclear whether exposure to the abnormal haemodynamic conditions associated with aortic stenosis leaves a persistent molecular and functional imprint on circulating monocytes, and how this haemodynamic imprint influences their subsequent differentiation and inflammatory behavior after their recruitment to the valvular extracellular matrix. In the present study, we demonstrate that pathological shear stress induces extensive proteomic and phosphoproteomic remodelling in human monocytes, leading to persistent alterations in signalling pathways that influence subsequent macrophage differentiation and lipid handling once the cells are incorporated into the model of valvular extracellular matrix. By integrating unbiased phosphoproteomics with pharmacological and genetic validation, we identified a previously unrecognized mechanotransducive pathway through which pathological shear stress induces a persistent Piezo1-dependent signalling imprint in monocytes, characterised by calcium influx, PKC activation and suppression of AKT signalling. Importantly, these haemodynamically primed monocytes subsequently exhibited altered inflammatory behavior within a bioengineered extracellular matrix model recapitulating a key pathological feature of CAVD, namely the accumulation of glycosaminoglycans such as chondroitin sulfate within the valve tissue ^36^. Together, these findings demonstrate that abnormal haemodynamics not only activate mechanosensitive signalling in circulating monocytes but also condition their subsequent responses within the diseased valve microenvironment. Our study therefore establishes a mechanistic link between pathological valve haemodynamics, extracellular matrix remodelling and chronic inflammation during CAVD progression.

Unlike endothelial cells and platelets, whose responses to disturbed flow have been extensively investigated, considerably less is known about how circulating monocytes respond to the extreme shear forces generated across stenotic aortic valves. Previous studies have largely viewed circulating monocytes as passive responders that become activated only after entering inflamed tissues. Our findings challenge this paradigm by demonstrating that pathological shear stress alone is sufficient to induce widespread molecular reprogramming before monocyte extravasation. Notably, these changes persisted following differentiation within a three-dimensional CS-enriched extracellular matrix, suggesting that the haemodynamic environment encountered in the circulation influences subsequent macrophage behaviour within the valve tissue. This concept of haemodynamic preconditioning provides a potential explanation for how mechanical forces contribute directly to the chronic inflammatory milieu characteristic of CAVD.

Our quantitative proteomic and phosphoproteomic analyses revealed rapid remodelling of signalling networks involved in calcium signalling, cytoskeletal organisation, cell adhesion and inflammatory regulation following exposure to pathological shear stress. These pathways are well recognised as central mediators of mechanotransduction and are essential for coordinating immune cell migration, adhesion and activation ^27, 37, 38^. Among the signalling pathways identified, kinase enrichment analysis consistently highlighted activation of PKC together with inhibition of AKT signalling, indicating that pathological shear stress induces coordinated mechanotransductive signalling rather than isolated phosphorylation events. PKC is a key regulator of monocyte activation and has been implicated in the production of pro-inflammatory cytokines ^38^. Consistent with these predictions, functional validation confirmed increased PKC activity following pathological shear stress, consistently circulating monocytes isolated from patients with severe CAVD exhibited elevated PKC activity prior to TAVI. Given the established role of PKC in promoting inflammatory activation, persistent PKC signalling may contribute to disease progression by priming circulating monocytes towards a hyper-inflammatory, pro-calcific phenotype before their recruitment into the valve^39^.

Piezo1 has emerged as a key mechanosensitive ion channel that enables immune cells to sense and respond to mechanical stimuli through calcium influx ^28^. Previous studies have demonstrated that Piezo1 regulates inflammatory activation of monocytes and macrophages ^6,21^ and mediates shear stress-induced neutrophil extra cellular trap formation ^7^. Consistent with these observations, our genetic studies using myeloid-specific Piezo1 knockout mice demonstrated that deletion of Piezo1 markedly attenuated Yoda1-induced calcium responses and significantly reduced calpain activation following pathological shear stress. These findings position Piezo1 upstream of the calcium-dependent signalling cascade identified in our phosphoproteomic analyses and establish Piezo1 as a critical mechanosensor linking pathological haemodynamic forces to intracellular signalling in monocytes.

PKC signalling has previously been implicated in mechanotransduction, inflammatory activation and regulation of macrophage function. Consistent with the kinase predictions generated from our phosphoproteomic analyses, pharmacological inhibition of PKC significantly altered oxLDL uptake and cholesterol transporter expression following haemodynamic conditioning. These findings suggest that PKC functions as an important downstream effector within the Piezo1-calcium-calpain signalling axis. Rather than acting as an isolated pathway, PKC appears to integrate calcium-dependent mechanotransduction with pathways regulating macrophage lipid homeostasis, providing a mechanistic explanation for how abnormal haemodynamic forces influence macrophage function within the diseased valve environment.

An interesting observation from the present study is that pathological shear stress altered macrophage lipid handling without necessarily promoting extensive foam cell formation. Although foam cells are a hallmark of atherosclerotic plaques, recent histological analyses have demonstrated that advanced calcified aortic valves contain relatively few foam cells despite evidence of lipid deposition during the early stages of disease. Instead, progressive CAVD is characterised predominantly by chronic inflammation, extracellular matrix remodelling and osteogenic differentiation ^40, 41^. Our findings are consistent with this evolving view and suggest that haemodynamic conditioning of circulating monocytes contributes primarily to inflammatory and phenotypic reprogramming rather than simply increasing lipid accumulation. Thus, the principal consequence of pathological shear stress may be to alter macrophage function within the valve microenvironment, thereby sustaining chronic inflammation that drives disease progression.

In conclusion, we propose a model in which pathological shear stress experienced by circulating monocytes during aortic stenosis initiates a Piezo1-dependent calcium signalling cascade that activates calpain and downstream PKC signalling, leading to persistent changes in macrophage differentiation and lipid handling after tissue recruitment. These findings identify a previously unrecognised mechanism by which abnormal haemodynamics directly regulate innate immune cell function and provide evidence that mechanical conditioning of circulating monocytes contributes to chronic inflammation in CAVD. Targeting components of this mechanotransductive signalling axis may therefore represent a novel therapeutic strategy to slow disease progression by interrupting inflammatory responses before irreversible valve remodelling and calcification occur.

